# Placental cytotrophoblast microvillar stabilization is required for cell-cell fusion

**DOI:** 10.1101/2024.02.22.581647

**Authors:** Wendy K. Duan, Sumaiyah Z. Shaha, Khushali J. Patel, Ivan K. Domingo, Meghan R. Riddell

## Abstract

The placenta is an essential organ of pregnancy required for maternal-fetal transport and communication. The surface of the placenta facing the maternal blood is formed by a single giant multinucleate cell: the syncytiotrophoblast. The syncytiotrophoblast is formed and maintained via fusion of progenitor cytotrophoblasts. Cell-cell fusion is a tightly regulated process, and in non-trophoblastic cells is accompanied by stereotypical alterations in cell shape by cells that have attained fusion-competence. The most prominent feature is the formation of actin-based membrane protrusions, but whether stereotypic morphological changes occur in fusion-competent cytotrophoblasts has not been characterized. Using a human placental explant model, we characterized cell shape factors associated with the attainment of cytotrophoblast fusion competence. We found that fusion-competent cytotrophoblasts are hypertrophic, elongated cells, that form microvilli at the apical membrane. The actin-membrane cross linker protein ezrin was found to have highly polarized expression within cytotrophoblast microvilli. Inhibition of ezrin activation destabilized cytotrophoblast microvilli and prevented cytotrophoblast fusion. Thus, we propose that the polarized activation of ezrin within apical microvilli and actin-mediated changes in membrane dynamics are necessary for cytotrophoblast fusion.

**Summary statement:** Fusion-competent cytotrophoblasts undergo dynamic changes in cell morphology including the acquisition of apically localized microvilli. Microvillar stabilization facilitates effective fusion and differentiation.

## Introduction

The placenta is an embryonically derived organ of pregnancy that carries out critical functions such as maternal-fetal nutrient exchange, hormone production, and protection from pathogens and maternal leukocytes. Trophoblasts are the placental epithelial lineage that comprise the maternal blood-facing placental surface. In humans, the syncytiotrophoblast (ST), a giant multinucleate single cell, covers nearly the entire maternal blood-facing surface. Therefore, this embryonically derived single cell is both a fetal sentinel and nutrient exchanger within the maternal compartment since it is bathed in maternal blood on its apical surface. By the end of gestation, the ST has been estimated to be ∼12m^2^ and contain billions of nuclei (Benirschke, 2000; Simpson et al., 1992). Importantly, the ST is post-mitotic and it is therefore maintained and expands via the incorporation of its underlying progenitor villous cytotrophoblasts (vCT) via cell-cell fusion. Adequate formation and renewal of the ST via vCT fusion is critical for proper placental function. Hence, altered vCT fusion is a feature of the common pregnancy complications intrauterine growth restriction (IUGR) and preeclampsia (Gauster et al., 2009; Langbein et al., 2008; Ruebner et al., 2010), which have a shared etiology of placental malformation. But much remains to be understood about the regulation of vCT fusion.

Cell-to-cell fusion (hereafter referred to as cell fusion) is a relatively uncommon but essential event in human biology. It occurs between gametes when the sperm and oocyte fuse, in myoblasts to maintain and form myofibers, and in the aforementioned fusion between cells of the villous trophoblast lineage. Fusion is a dramatic cellular event that is highly regulated to ensure it occurs between the right cells at the correct location within a tissue, and requires the expression of fusogens, proteins that catalyze the local disruption of the lipid bilayers and their rejoining. It has been proposed that fusion can be broken down into three stages: 1): attainment of fusion competence; 2) commitment to fusing; 3) the fusion event (Aguilar et al., 2013). In myoblasts, these stages have been extensively characterized in both mammalian and *Drosophila* cells (Abmayr and Pavlath, 2012; Kim and Chen, 2019; Petrany and Millay, 2019), but our understanding of the regulation and execution of trophoblast fusion is less developed.

Single cell-RNA-seq and single nuclei-seq analyses have revealed that multiple transcriptionally distinct populations of vCT exist at one time in the first trimester placenta (Arutyunyan et al., 2023; Shannon et al., 2022; Wang et al., 2024). These analyses have identified vCT with higher expression of ST marker genes that are assumed to be a population that has attained fusion competence due to high expression of the placental fusogen syncytin-2 (*ERVFRD-1*) (Arutyunyan et al., 2023; Shannon et al., 2022; Wang et al., 2024). Similarly, ultrastructural examination of first trimester and term placenta has revealed a sub-population of vCT that more strongly resemble the ST in their nuclear structure and organellular composition (Jones and Fox, 1991). It is well established that commitment to fusion is accompanied by changes in cell shape. For example, myoblasts are known to change their cell shape to laterally align with the myofiber with which they will fuse (Lehka and Redowicz, 2020), and the appearance of membrane protrusions on the cell surface of fusion competent cells is also a highly conserved feature amongst fusogenic cell types (Brukman et al., 2019; Wilson and Snell, 1998). These shape changes appear to have the purpose of bringing fusing membranes into closer approximation (Brukman et al., 2019; Kim and Chen, 2019; Wilson and Snell, 1998), and alterations in cell surface area immediately after fusion has been proposed to regulate transcriptional commitment to a differentiated state via an AMPK-YAP1 signaling pathway (Feliciano et al., 2021). Bundled filamentous-actin (F-actin) is a core cytoskeletal component of membrane protrusions observed in fusion competent myoblasts where the formation of invasive membrane podosomes allows for efficient fusogen engagement and mechanical coordination between the membranes of apposing cells (Kim and Chen, 2019; Kim et al., 2015; Petrany and Millay, 2019; Sens et al., 2010; Shilagardi et al., 2013). Membrane projections have been observed by brightfield live cell imaging between fusing BeWo trophoblastic choriocarcinoma cells (Wang et al., 2014) and the variable presence of membrane protrusions in primary vCT undergoing spontaneous fusion has been observed by scanning electron microscopy (SEM) (Bax et al., 1989), but the cytoskeletal components driving these protrusions and characterization of other shape factors that may distinguish fusion competent vCT have not been identified.

The actin-membrane linker protein ezrin has been shown to play an important, though complex, role in the formation and stabilization of membrane protrusions (Saotome et al., 2004; Welf et al., 2020) and is particularly important for the maintenance of more stable membrane structures like epithelial microvilli (Pelaseyed and Bretscher, 2018). Ezrin requires activation via phosphorylation on its Thr-567 residue in order to attain an open conformation and bind both the F-actin filament and overlying membrane (Zhu et al., 2007). Interestingly, ezrin has been shown to function as an A-kinase anchoring protein (AKAP) in fusing 2D primary vCT cultures and serves as a scaffolding protein to protein kinase A (PKA), an important regulator of vCT fusogen expression (Gerbaud et al., 2015; Pidoux et al., 2014) but whether it also functions as a regulator of vCT shape alterations to enable fusion remains to be examined.

Here, we characterized cellular shape parameters of the vCT layer in intact human first trimester placental tissue. To identify the shape characteristics that are associated with fusion commitment, we exploited a first trimester explant model where widespread spontaneous vCT fusion occurs after the removal of the overlying ST layer (Baczyk et al., 2009; Baczyk et al., 2013; Miller et al., 2005). We identified that the population of vCTs stimulated to fuse by the removal of the ST are enriched in cells that accumulate surface area in the apical region and produce F-actin and ezrin containing microvilli. Chemical inhibition of ezrin activation in ST-stripped explants strongly abrogated vCT fusion, altered the pattern of F-actin on the apical surface, and reduced expression of ST marker genes. Therefore, we have uncovered that vCT fusion requires the polarized accumulation and activation of ezrin within apically localized microvilli.

## Results

### ST regeneration explant model recapitulates syncytialization *in vitro*

To establish a model that would allow for the accumulation of fusion-competent vCT, we used a first trimester human placental explant model. Tissue was briefly trypsinized to promote ST denudation and then cultured to allow for ST shedding and regeneration. Unlike established two-dimensional trophoblast stem cell culture (Okae et al., 2018), primary cultured vCT (Pidoux et al., 2014), and organoid models (Haider et al., 2018; Sheridan et al., 2020; Turco et al., 2018) this model allows for the observation of three dimensional spontaneous syncytialization from pre-existing vCT subpopulations on an intact basement membrane. Near complete removal of the pre-existing ST was achieved by 24hrs post-trypsinization (Fig. 1A), exposing a mononucleate anti-E-cadherin and anti-integrin-α-6 positive vCT layer (Aghababaei et al., 2015; Aplin, 1993; Longtine et al., 2012). By 72hrs post-trypsinization, a multinucleated structure displaying clear anti-β-human chorionic gonadotropin (β-hCG) signal (ST marker) atop or interspersed with anti-integrin-⍺-6-positive or anti-E-cadherin-positive mononucleated vCTs was consistently observed (Fig. 1A). The formation of multinucleated structures often led to the exhaustion of the vCT layer, resulting in regions of ST without underlying vCTs; however, cases where vCTs were present underneath a regenerated ST were also observed. The degree of ST regeneration was quantified by a significant increase in the relative ST:vCT coverage between 24hrs and 72hrs post-trypsinization (Fig. 1B). Thus, this model enriches vCT into a fusion competent state to allow for widespread synchronized ST regeneration.

**Figure 1:**
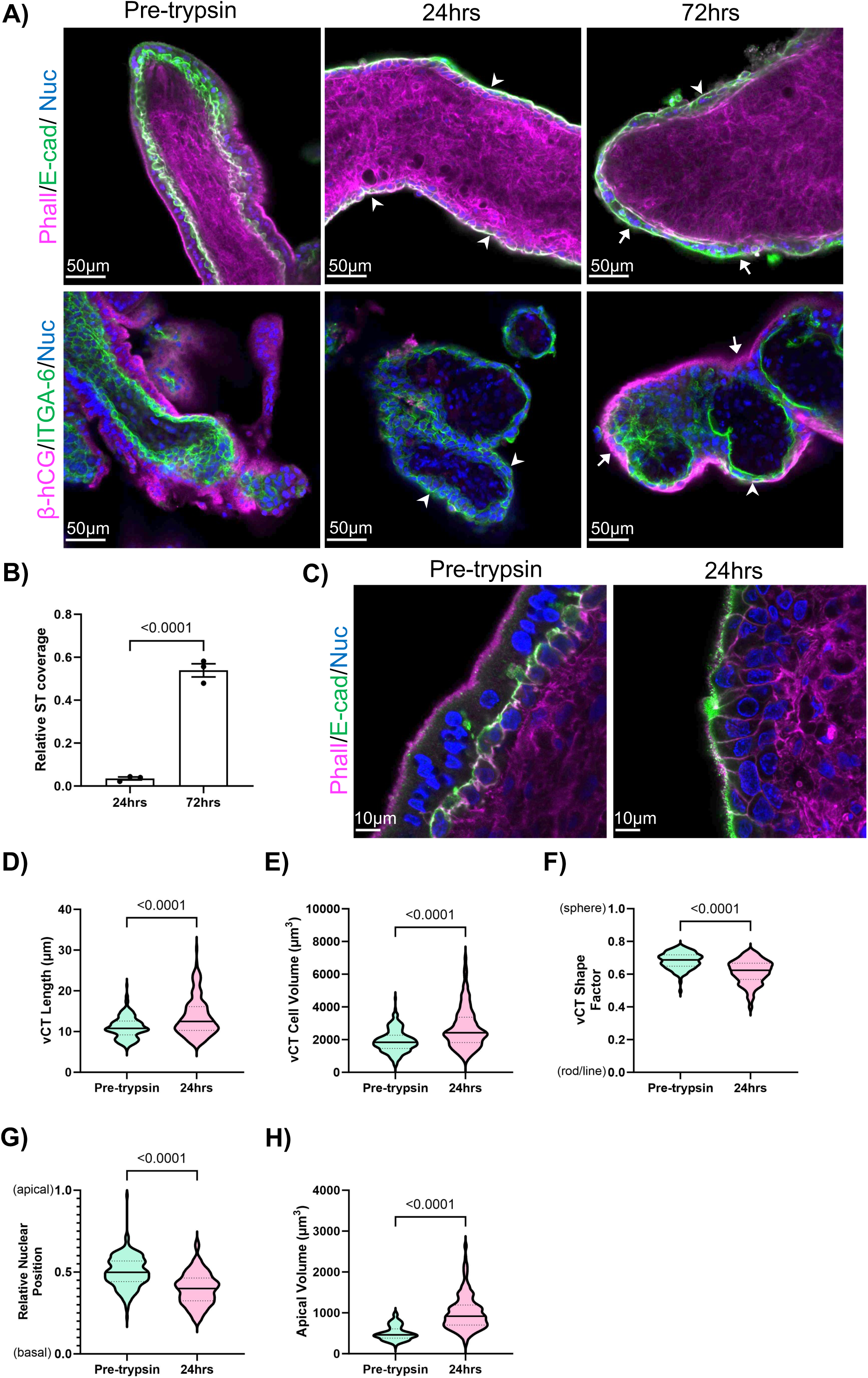
ST regeneration model enriches a fusion-competent vCT subpopulation and synchronizes ST development. A) Representative single XY-plane confocal microscopy images of placental tissue in pre-trypsinized non-cultured controls, at 24hrs post-trypsinization, and 72hrs post-trypsinization; top row was stained with phalloidin (F-actin; magenta), anti-E-cadherin (green), and Hoechst 33342 (nuclei; blue); bottom row was stained with anti-β-hCG (magenta), anti-integrin-⍺-6 (green), Hoechst 33342 (nuclei; blue); white arrows indicate ST regions; arrowheads indicate vCT regions; B) Summary data of ST coverage (ST:vCT) at 24hrs and 72hrs normalized to pre-trypsin controls; mean ± S.E.M., unpaired parametric student’s t-test, *n=*3; ST coverage (ST:vCT) was calculated using equation 2 (Methods); C) Representative single XY-plane confocal microscopy images of pre-trypsinized control tissue and tissue 24hrs post-trypsinization stained with phalloidin (magenta), anti-E-cadherin (green), and Hoechst 33342 (nuclei; blue); D) Summary data of vCT cell length (µm); *n_pretrypsin_=*210; *n_24hrs_=*178; E) Summary data of vCT cell volume (µm^3^); *n_pretrypsin_=*237; *n_24hrs_=*290; F) Summary data of vCT shape factor; *n_pretrypsin_=*237; *n_24hrs_=*290; G) Summary data for the relative position of the nucleus along the apical-basal axis; *n_pretrypsin_=*210; *n_24hrs_=*178; calculated using equation 1 (Methods); H) Summary data of vCT apical volume (µm^3^); *n_pretrypsin_=*195; *n_24hrs_=*220; D) - H) unpaired Mann-Whitney test; bold line = median; dashed lines = quartile 1 and quartile 3.

### ST denudation leads to enrichment of a morphologically distinct vCT subpopulation

Since the removal of the ST layer led to depletion of the vCT layer and regeneration of a multinucleate ST, we surmised that there was a time-dependent enrichment in fusion competent vCT, allowing for the identification of their characteristic morphology. Therefore, we acquired high magnification z-stack confocal images of uncultured placental tissue and cultured explants at 24hrs post-trypsinization from the same donor placentas after whole-mount staining with anti-E-cadherin, phalloidin, and a nuclear dye (Fig. 1C) to quantify population-wide shifts in morphological parameters observed with ST denudation. ST-denuded explants were enriched with hypertrophic vCTs that resulted in an upward shift in the median cell length (Fig. 1D) and median cell volume (Fig. 1E) compared to ST-intact placental tissue. Accompanying this growth, ST-denuded vCT populations shifted towards an elongated and less spherical cell shape, resulting in a downward shift in shape factor (Fig. 1F). The median position of the vCT nucleus in ST-intact placental tissue was found to be halfway between the apical and basal membrane; however, the median nuclear position in ST-denuded vCTs shifted closer to the basal membrane (Fig. 1G). On the apical surface, ST-denuded vCTs were found to have significantly increased apical volumes compared to ST-intact vCTs (Fig.1H). Thus, ST denudation pushes vCTs to undergo morphological changes including changes in cell size, shape, and polarity, thereby enriching a structurally distinct vCT subpopulation prior to fusion.

### Microvilli decorate the vCT apical surface prior to fusion *in vitro* and *in vivo*

One of the most striking and highly enriched features of vCT in denuded explants was an increase in apical volume that was mostly attributable to an increased occurrence of F-actin-positive hair-like projections at the apical surface (Fig. 1C, 2A). Bundled F-actin in fusion-competent cells has been established as central to membrane protrusion formation (Kim and Chen, 2019; Kim et al., 2015; Petrany and Millay, 2019; Sens et al., 2010; Shilagardi et al., 2013), therefore we hypothesized that membrane projections were accumulating on the apical surface of vCT in our explant model. Scanning electron micrographs (SEM) of the vCT surface at 24hrs post-trypsinization revealed varying amounts of membranous protrusions ∼100nm in diameter on the apical surface of individual vCTs and bridging the intercellular gaps between individual cells (Fig. 2B). A wide variation in the amount of these structures on individual cells was observed within these micrographs. Most vCTs were moderately to heavily decorated on their apical surface, although vCTs devoid of these structures were also observed. Berryman *et al*. previously reported immunogold labelling of ezrin, a core component of epithelial microvilli, within apical membrane projections of sporadic vCT in intact placenta of an unreported gestational age using transmission electron microscopy (TEM) (Berryman et al., 1993). Therefore, we stained ST-denuded explants with an anti-ezrin antibody to confirm whether a similar signal pattern of apically accumulated ezrin existed in our model. As expected, anti-ezrin signal strongly localized to the apical surface of both mononucleated vCTs and fused binucleated regenerated STs (Fig. 2C). Based on the relative diameter of the apical membrane structures and the presence of F-actin and ezrin within them we concluded that these membrane structures are microvilli as opposed to the invasive podosomes characteristic of fusion competent myocytes. Altogether, these data suggest that the formation of extensive microvilli on the apical surface of the vCT surface represents a morphological adaptation associated with a fusion competent state.

**Figure 2:**
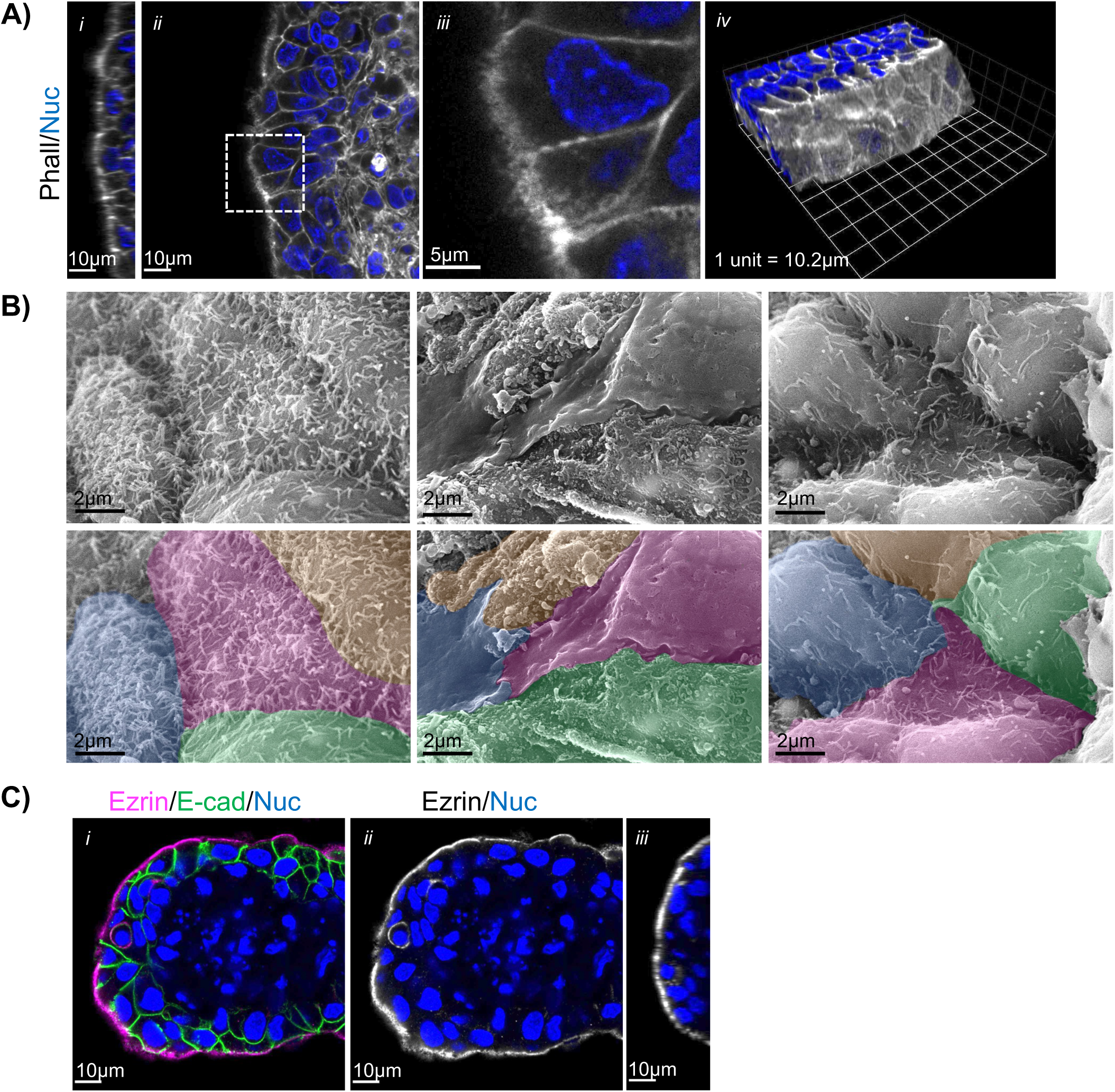
ST denudation increases accumulation of microvilli on the vCT apical membrane. A) Representative single ZY-plane (*i)*, XY-plane (*ii, iii*), and 3D reconstituted (*iv*) confocal microscopy images of tissue 24hrs post-trypsinization; merged image of phalloidin (greyscale) and Hoechst 33342 (nuclei; blue) signals; *iii* = higher magnification image of indicated region in *ii*; B) Representative SEM images of explant tissue 24hrs post-trypsinization; bottom panels = false-coloured images with colours marking individual vCT; C) Representative single XY-plane confocal microscopy images (*i, ii*) of tissue 24hrs post-trypsinization; *i* = merged image of anti-ezrin (magenta), anti-E-cadherin (green), and Hoechst 33342 (nuclei; blue) signals; *ii* = isolated anti-ezrin (greyscale) and Hoechst 33342 (nuclei; blue) signals; *iii*= ZY plane isolated anti-ezrin (greyscale) and Hoechst 33342 (nuclei; blue) signals.

We then sought to confirm the observations of Berryman *et al*. to identify whether vCT with apical F-actin and ezrin containing structures could be observed in intact first trimester placenta. Microvilliation of the vCT apical (ST-facing) membrane of a subset of cells has also been observed in additional TEM studies of both term and first trimester placenta (Jones and Fox, 1991; Tashev et al., 2022). As expected, we observed vCT with apical membrane projections at the vCT-ST interface with approximately the same diameter of those observed on the apical surface of vCT in our denuded explant model in TEM micrographs (Fig. 3A), though extensive microvilli were not clearly visible in the majority of vCT, aligning with published data (Jones and Fox, 1991; Tashev et al., 2022). Confocal z-stack images of whole-mount intact placental tissue revealed that individual vCT possess varying degrees of apical hair-like F-actin (phalloidin) signal (Fig. 3B), similar to the ST regeneration model. Interestingly, towards the tip of individual villi there was a visible increase in the occurrence of highly branched and fine apical phalloidin signal in vCT while along the length of the villus vCT more often displayed a largely smooth, unbranched apical phalloidin staining pattern.

**Figure 3:**
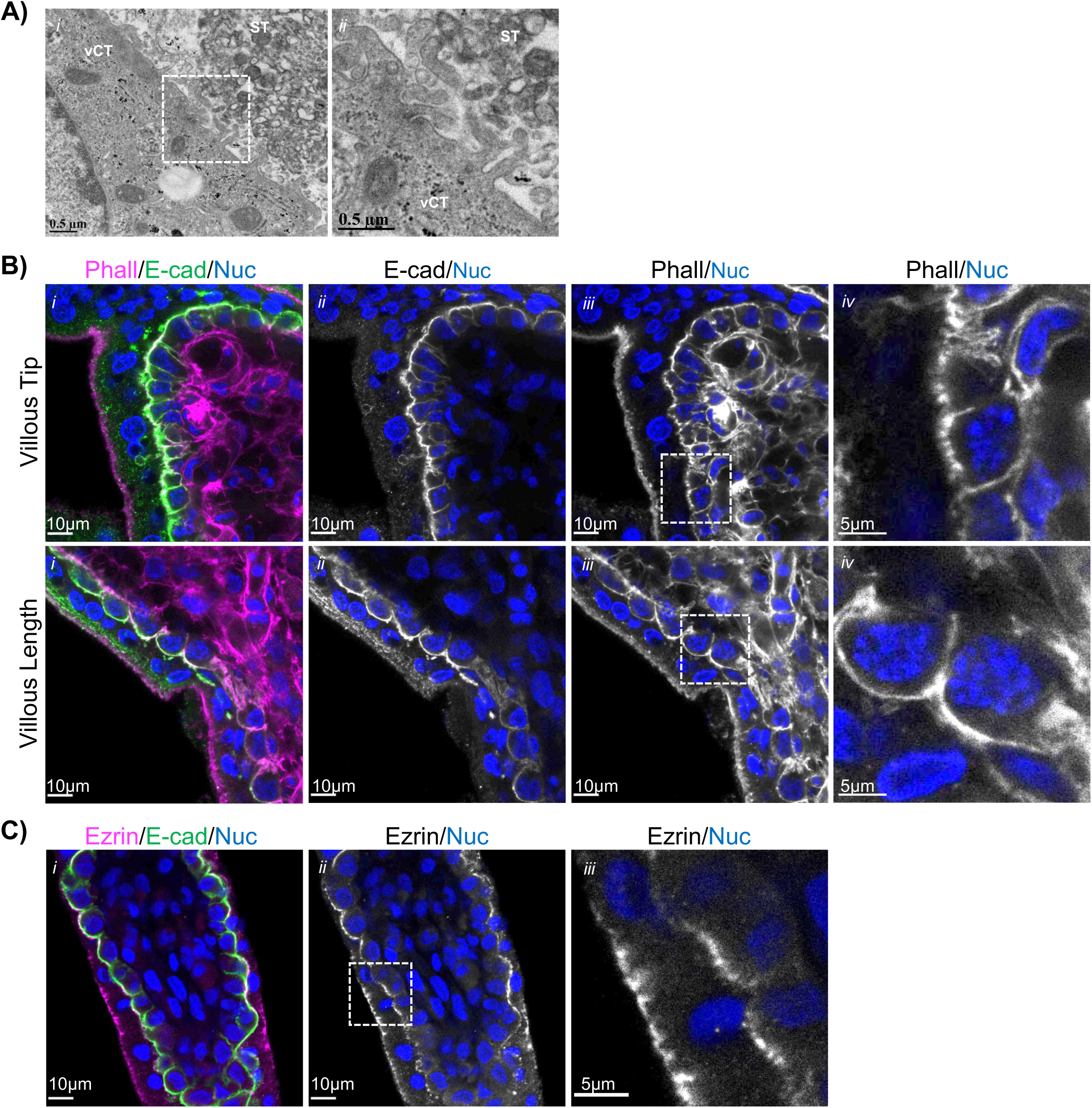
Apical microvilli of are present in vCT in intact placental tissue. A) Representative TEM images of uncultured placental tissue; *ii* = higher magnification image of indicated region in *i*. B) Representative single XY-plane confocal microscopy images of uncultured placental tissue; *i* = merged images of phalloidin (magenta), anti-E-cadherin (green), and Hoechst 33342 (nuclei; blue) signals; *ii* = isolated anti-E-cadherin (greyscale) and Hoechst 33342 (nuclei; blue) signals; *iii* = isolated phalloidin (greyscale) and Hoechst 33342 (nuclei; blue) signals; *iv* = higher magnification images of indicated regions of isolated phalloidin (greyscale) and Hoechst 33342 (nuclei; blue) signals; C) Representative single XY-plane confocal microscopy images of uncultured placental tissue; *i* = merged images of anti-ezrin (magenta), anti-E-cadherin (green), and Hoechst 33342 (nuclei; blue); *ii* = isolated anti-ezrin (greyscale) and Hoechst 33342 signals; *iii* = higher magnification image of indicated region in *ii*.

Consistent with the findings in our *in vitro* model, anti-ezrin signal variably localized to the apical surface of individual vCT and, as previously reported, in a discontinuous pattern at the apical surface of the ST (Fig. 3C) (Berryman et al., 1993; Patel et al., 2023). Therefore, we confirmed that apical microvilli are present in a subset of vCT in the first trimester placenta. In combination with our data showing the accumulation of cells with this feature in denuded placental explants it suggests that the assembly of apical microvilli is a previously unrecognized stage in fusion commitment for vCT.

### Microvillar stabilization via activated ezrin is needed for vCT fusion and differentiation

Since vCT microvilliation is a conserved feature *in vivo* and *in vitro*, we hypothesized that formation of apical microvilli may be obligate for trophoblast fusion. To target microvilli without disrupting whole cell actin dynamics, we selected to treat vCT in our ST-regeneration explant model with NSC668394, which binds directly to ezrin to prevent its phosphorylation at the Thr-567 residue that is necessary for ezrin activation, membrane binding, and microvilli stabilization (Viswanatha et al., 2012; Welf et al., 2020). When ST-denuded explants were pulsed with ezrin inhibitor for 2hrs, there was a smoothening of the actin signal at the apical membrane and a loss of the hair-like F-actin projections (Fig. 4A), showing that disruption of ezrin activation leads to the destabilization of apical microvilli. In addition, ezrin inhibitor dose-dependently impaired vCT fusion when explants were treated for 48hrs (Fig. 4B, C). While exposure to 5µM and 10µM of inhibitor had variable effects on fusion, 20µM and 50µM doses resulted in a consistent 63% and 86% decrease in fusion respectively when compared to vehicle controls. Therefore, blocking ezrin activation strongly inhibits vCT fusion and leads to a loss of apical microvilli.

**Figure 4:**
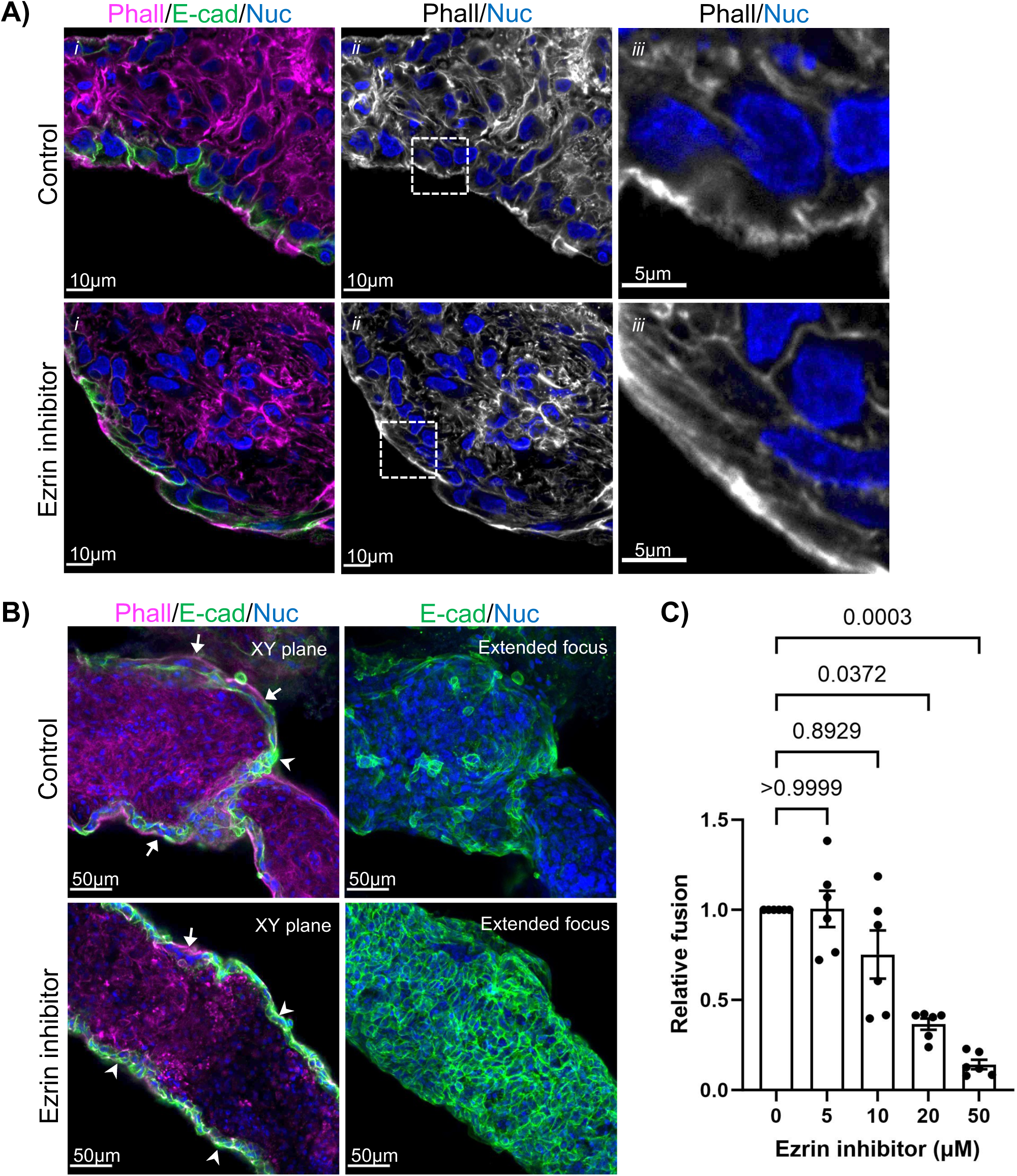
Ezrin inhibitor treatment impairs vCT fusion and alters protrusive actin filament accumulation on the apical surface. A) Representative single XY-plane confocal microscopy images of explants pulsed for 2hrs ± 50µM ezrin inhibitor at 24hrs post-trypsinization; *i* = merged image of phalloidin (magenta), anti-E-cadherin (green), and Hoechst 33342 (nuclei; blue) signals; *ii* = isolated phalloidin (greyscale) and Hoechst 33342 (nuclei; blue) signals; *iii* = higher magnification image of regions indicated in *ii*; B) Representative single XY-plane and extended focus confocal microscopy images of explants cultured for 72hrs total ± 50µM ezrin inhibitor added at 24hrs post-trypsinization stained with phalloidin (magenta), anti-E-cadherin (green), and Hoechst 33342 (nuclei; blue); white arrows indicate regenerated ST regions; arrowheads indicate unfused vCT regions; C) Summary data of dose-dependent ezrin inhibitor treatment on fusion (change in ST:vCT) normalized to vehicle controls; mean ± S.E.M., unpaired Kruskal-Wallis test with Dunnett’s multiple comparisons, *n=*6; fusion (change in ST:vCT) is calculated using equation 3 (Methods).

Cell shape, or more specifically, a decrease in cell surface area immediately after fusion, has recently been shown to reinforce transcriptional regulation of cell differentiation in fused cells (Feliciano et al., 2021). Multiple studies examining trophoblast fusion and differentiation using the BeWo choriocarcinoma cell line and primary vCT have identified that trophoblast fusion and functional differentiation into ST, or at least the production of β-hCG, are isolatable processes (Collett et al., 2012; Johnstone et al., 2005; Orendi et al., 2010). Therefore, we questioned whether ezrin inhibitor treatment would also block β-hCG expression and the expression of other vCT differentiation and ST marker genes. Ezrin inhibitor treatment led to an almost a complete loss of anti-β-hCG signal compared to vehicle controls (Fig. 5A), and a concomitant decrease in *CGB* mRNA (Fig. 5B). Additional markers of vCT differentiation and ST were also assessed by RT-PCR. *GCM1* is present nearly exclusively in the human placenta (Baczyk et al., 2013; Nait-Oumesmar et al., 2000) and is a key placental transcription factor for vCT differentiation into ST (Baczyk et al., 2009; Jeyarajah et al., 2022; Woods et al., 2018), and its levels increase as syncytialization progresses (Jeyarajah et al., 2022). With ezrin inhibition, *GCM1* expression in explant lysates was significantly decreased compared to paired controls (Fig. 5C). Syncytin-1 (*ERVW-1*) and syncytin-2 (*ERVFRD-1*) are vCT fusogens (Malassine et al., 2007; Mi et al., 2000; Muir et al., 2006) and therefore essential for mediating vCT fusion(Blond et al., 2000; Frendo et al., 2003; Mi et al., 2000; Vargas et al., 2009). A significant increase in relative *ERVFRD-1* expression (Fig. 5D) and a non-significant increase in *ERVW-1* abundance (Fig. 5E) was observed with ezrin inhibition, indicating that blocking ezrin activation stalled vCT differentiation into ST. When taken together with the significant decrease in cell fusion observed in Fig. 4, these data suggest a loss of fusogen efficacy, not expression, occurs in the absence of ezrin activation. In summary, ezrin activity and stabilization of microvillar structure is necessary for fusion and loss of ezrin activity effectively arrests syncytialization.

**Figure 5:**
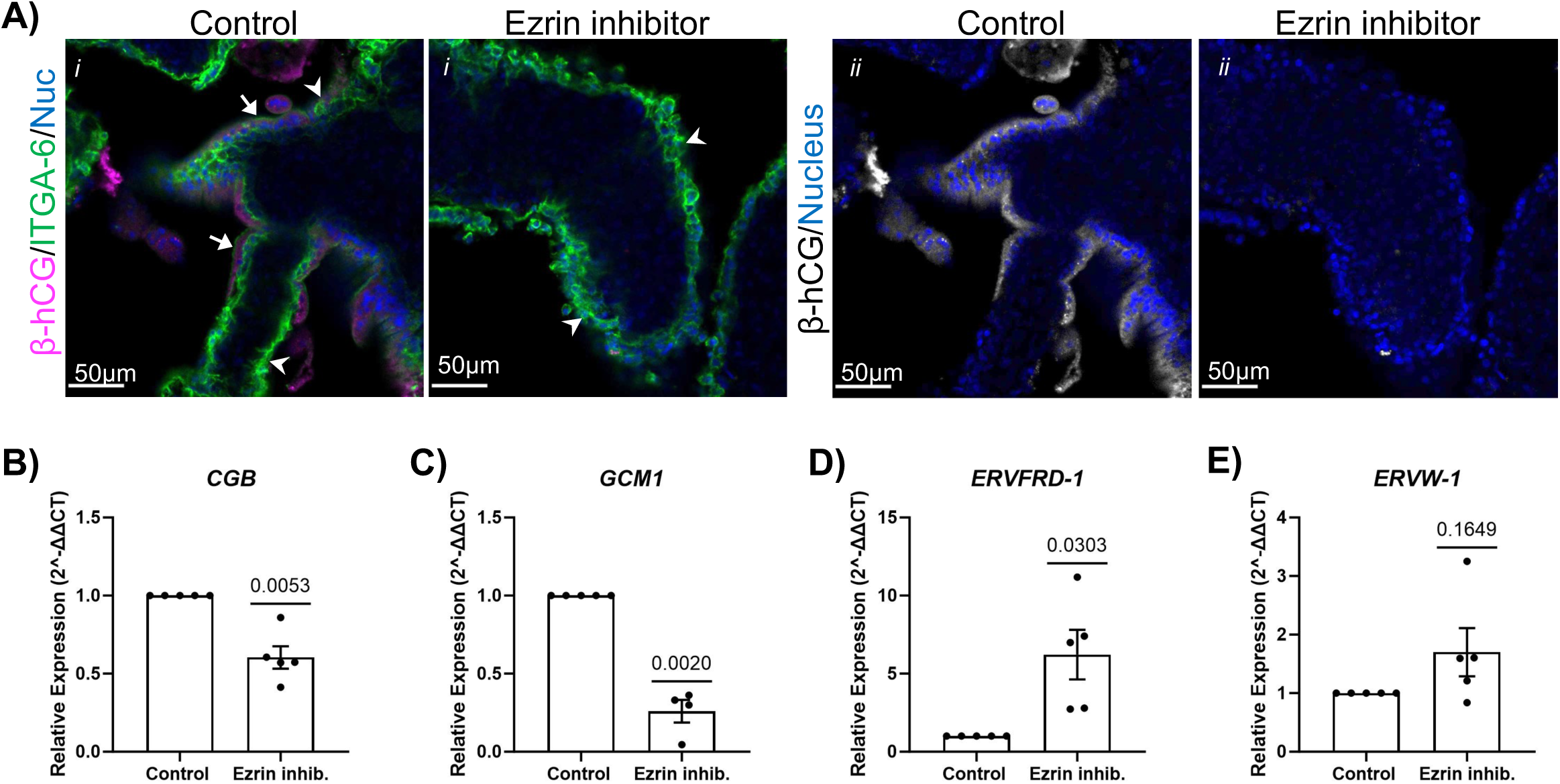
Ezrin inhibitor treatment impairs vCT functional differentiation. A) Representative single XY-plane confocal microscopy images of explants cultured for 48hrs ± 50µM ezrin inhibitor added 24hrs post-trypsinization; *i* = merged images of anti-β-hCG (magenta), anti-integrin-⍺-6 (green), and Hoechst 33342 (nuclei; blue) signals; *ii* = isolated anti-β-hCG (greyscale) and Hoechst 33342 (nuclei; blue) signals; white arrows indicate regenerated ST regions; arrowheads indicate unfused vCT regions; B) Relative *CGB* expression C) Relative *GCM1* expression D) Relative *ERVFRD-*1 expression, and E) Relative *ERVW-1* expression in explants cultured for 48hrs total ± 50µM ezrin inhibitor added 24hrs post-trypsinization; Data represented as 2^-ΔΔCT to mean of *TOP1* and *CYC1* normalized to controls; mean ± S.E.M., one-sample t-test, *n*=4-5.

## Discussion

Proper formation and maintenance of the ST throughout gestation is critical for placental function and significantly contributes to healthy pregnancy progression. Diminished vCT fusion capacity is a feature of devastating pregnancy complications like IUGR and preeclampsia (Gauster et al., 2009; Langbein et al., 2008; Ruebner et al., 2010), which are associated with increased perinatal morbidity and mortality and lifelong increased risk of cardiovascular and neurological disease for affected infants (Burton et al., 2016). Therefore, thorough understanding of the molecular regulation of vCT fusion could allow for the development of treatments for harmful placental pathologies and improve lifelong health. Here, we found that the stabilization of microvillar membrane protrusions within vCT is necessary for successful fusion between neighbouring cells for ST regeneration. We also present data that suggests that stabilized vCT microvilli may be necessary for fusion with intact ST, thereby confirming that trophoblasts require protrusive F-actin-based membrane structures to successfully catalyse cell-cell fusion as has been seen in all other fusing cell types.

Our data shows that the membrane binding protein ezrin plays a critical role in regulating vCT membrane structure and fusion. Importantly, ezrin has previously been shown to regulate vCT fusion through its role as a scaffolding protein for PKA and the gap junction protein connexin-43 (Pidoux et al., 2014). PKA phosphorylation of connexin-43 allows for the formation of gap junctions between adjacent cells to facilitate signal synchronization before fusion occurs. Using primary 2D vCT culture and ezrin Thr-567 mutant overexpression, Pidoux *et al*. have shown that the active form of ezrin is required for it to serve as a PKA scaffold. Therefore, we cannot rule out that the significant effects we observed with ezrin inhibitor treatment on fusion are not at least partially due to disruption in PKA/connexin-43 binding and our results are unlikely to be due to the loss of microvillar structures alone. Irrespective of the relative contribution of microvillar structure versus PKA scaffolding on fusion, the combination of our analyses and this past work highlight that vCT accumulation of ezrin within microvillar structures localizes critical pro-fusogenic signaling within a highly specialized cellular compartment within the apical region of the cell. Intestinal microvilli are known to contain their own sub-proteome (McConnell et al., 2011) and the polarized localization of proteins within highly specialized structures may be driven by the differential composition of the microvillar lipid bilayer within highly curved and tightly packed structures (Cebecauer, 2021; Ikenouchi et al., 2013). The concept of microvilli as signaling centres is established in diverse non-fusing cell types like intestinal epithelia and T-cells (Kim et al., 2018; McConnell et al., 2009). Therefore, stabilized microvilli within the vCT seem likely to serve as a signaling platform to enrich the necessary components for fusion in proximity to the overlying ST and warrant further investigation.

In addition, other fusing cell systems have shown the importance of increasing membrane-membrane contact sites for fusion (Brukman et al., 2019; Kim and Chen, 2019; Wilson and Snell, 1998). Therefore, microvilli may serve to increase the apical surface area of vCT for this purpose. E-cadherin signal was strongly accumulated in proximity to the observed apical ezrin signal in our experiments, and others have observed the gap junction protein connexin-43 in a similar pattern (Cronier et al., 2002), thus microvilli may facilitate increased contact points by concentrating junctional protein interactions between the vCT and overlying ST. Simply due to the proximity of vCT microvilli to the ST membrane, it is tempting to suggest that fusion pores may also arise in microvillar tips. Syncytin-2 has been shown to have a non-polarized membrane localization in select vCT *in vivo* (Esnault et al., 2008), but as a class I fusogen (Vance and Lee, 2020), it requires engagement with its receptor major facilitator superfamily domain containing 2 (MFSD2) (Esnault et al., 2008) allowing for its extended conformation and subsequent insertion into the adjacent cell membrane to facilitate formation of a fusion pore. Therefore, microvilliation may allow for more efficient syncytin-2/MFSD2 engagement due to the diminished space between vCT and ST membranes and increased contact sites. MFSD2 is expressed on both the ST and CT (Esnault et al., 2008), therefore increasing the contact points between vCT syncytin-2 and ST MFSD2 would increase the likelihood of fusion. Our ST denuded explant model takes advantage of the tendency of vCT to spontaneously fuse when overlying syncytium is lost. Though not a perfect physiologic system, regions of localized ST denudation and subsequent fibrin deposition are a feature of both healthy and pathologic placentas (Scifres and Nelson, 2009), therefore this model more accurately represents vCT fusion for ST regeneration than ST maintenance. Importantly, though our cell shape factor analyses revealed that ST denudation shifts vCT population medians for the analysed factors by increasing the proportion of cells that lie on the higher or lower extremes of a population, the majority of vCT in the denuded tissue still fall within the limits of those observed in intact tissue. Thus, the cellular features associated with fusion competence are likely accentuated by a lack of negative feedback from the ST.

Interestingly, the tendency for vCT to increase in length in the apical-basal axis, the positioning of the nucleus in a more basal position with ST removal, and the polarized accumulation of ezrin at the apical surface support a model whereby the initiation of an apical-basal polarization network amplifies a pro-fusogenic vCT state and dictates the position of the newly formed ST. Ezrin localization in 2D cultured primary vCT was not found to be polarized and generally localized throughout the membrane (Pidoux et al., 2014), suggesting that vCT interactions with the extracellular matrix of the basement membrane may allow for polarization, and thereby enhance fusion. Critically, apparent microvillar projections have been observed by others during 2D vCT fusion (Bax et al., 1989), indicating that polarization of ezrin within the cell is not obligate for vCT fusion, but supporting that the formation of microvilli likely is. Work with trophoblast organoids indirectly supports our hypothesis that vCT polarity and extracellular matrix interactions direct the position of ST formation. In Matrigel embedded organoids where the extracellular matrix is surrounding a ball of progenitor CTs, CTs spontaneously form ST in a cystic core instead of on the outer surface (Haider et al., 2018; Turco et al., 2018). Our previous work has shown that the canonical apical-basal polarity regulating factor atypical protein kinase-c (aPKC) does not have a highly polarized staining pattern within first trimester vCT (Shaha et al., 2022), suggesting an alternative pathway may be activated to reinforce vCT polarity during differentiation. Understanding the factors that initiate vCT polarity and microvillus formation will be an interesting future direction.

We chose to perform our analyses in first trimester tissue explants due to the continuous nature of the vCT layer at this point in gestation, thereby allowing us to analyze the greatest number of cells. With advancing gestation vCT change from a continuous layer of cuboidal cells to a discontinuous layer of flattened cells (Jones and Fox, 1991). Microvilliated vCT in term tissue have been observed in intact term placental sections with TEM (Jones and Fox, 1991), supporting that this may be a conserved feature of vCT fusion competence across gestation. Other shape parameters we examined may not translate to later gestational stages. Future work is necessary to explore whether fusion competent vCT beyond the first trimester have shared morphological features with first trimester cells. Differences in the way vCT execute fusion across gestation are possible and understanding their conserved and divergent regulation would inform development of treatments in the future.

In summary, we show that stabilization of vCT microvilli is critical for fusion and formation of ST. The microvillus is a conserved site of fusion throughout evolution seen in cell-fusion events in species as diverse as sea urchin and *Drosophila*, yet understanding of why these structures are necessary and how their emergence and stabilization is regulated to control one of the most dramatic events in cell biology is lacking. We identified that ezrin and F-actin are contained in vCT membrane protrusions when fusion is stimulated, therefore future work to address the temporal and spatial regulation of the cortical actin cytoskeleton and its interplay with ezrin and membrane proteins to regulate microvilli emergence and localization will be particularly important.

## Materials and Methods

### Tissue collection

9-12 weeks gestational age human placental samples were collected by methods approved by the University of Alberta Human Research Ethics Board (Pro00089293). First trimester placental tissue was obtained from elective pregnancy terminations following informed consent from the patients.

### ST regeneration model and treatments

Placental samples were cut into 2-3mm^3^ sized explants in ice-cold phosphate-buffered saline (PBS) and then either fixed immediately as uncultured pre-trypsin controls or trypsinized and cultured. For trypsinization, tissue was digested with 0.25% trypsin-EDTA (Gibco; Burlington, ON, Canada; cat:25200-072) for seven minutes at 37°C. The reaction was quenched using 10% heat-inactivated fetal bovine serum (FBS, Wisent; Saint-Jean-Baptiste, QC, Canada; cat:098150), and a single explant was placed in each well of a 48 well plate in medium consisting of Iscove’s Modified Dulbecco’s Medium (IMDM) (Gibco, cat:12440-053), 5% heat-inactivated FBS (Wisent, cat:098150), 50µg/mL gentamicin (Gibco, cat:15710-064), and 1X ITS-X supplement (Gibco, cat:51500-056) at 37°C 5% CO_2_. Medium was changed 24hrs post-trypsinization to remove stripped ST debris. For ezrin inhibitor treatments, 5-50µM, of NSC668394 (EMD Millipore; Oakville, ON, Canada; cat:341216, lot:3934373) or vehicle control (DMSO) was added to medium 24hrs post-trypsinization with 3-12 explants per treatment group for 2-48hrs.

### Tissue staining

Placental tissue (uncultured and cultured explants) was fixed using 2% paraformaldehyde (PFA) overnight at 4°C or 4% PFA on ice for two hours. When necessary, heat-induced antigen retrieval was conducted using 0.01M sodium citrate buffer at pH 6.0. Fixed tissue was blocked and permeabilized in blocking buffer (5% normal donkey serum, 0.5% Triton-X100). Tissue was then incubated overnight at 4°C in blocking buffer with anti-E-cadherin (R&D Systems; Minneapolis, MN, USA; cat:MAB1838), anti-integrin-⍺-6 (Stem Cell Technologies; Vancouver, BC, Canada; cat:60037), anti-β-human-chorionic-gonadotrophin (β-hCG) (Abcam; Cambridge, UK; cat:ab9582), and anti-Ezrin (Invitrogen; Burlington, ON, Canada; cat: PA5-18541). For further antibody information, please see Appendix Table 1. Tissue was then washed with PBS and 0.05% Tween (PBST) and PBS, and incubated with the appropriate fluorescent secondary antibodies AlexaFluor 488 (Invitrogen), AlexaFluor 594 (Invitrogen), and/or stained with 0.17µM phalloidin-iFluor 594 (AAT Bioquest; Pleasanton, CA, USA; cat:23122). Tissue was then incubated with 10µg/mL Hoechst 33342 (Thermo Fisher Scientific; Burlington, ON, Canada; cat:H3570) followed by PBS washes and whole-mounted with imaging spacers and Fluoromount-G mounting medium (Southern Biotech; Burlington, ON, Canada; cat: 0100-01).

### Confocal microscopy image capture and analyses

Triplicate z-stack images were acquired with a Zeiss LSM 700 confocal microscope using a Zeiss Plan Apochromat-20X/0.8 M27 or Zeiss Plan Apochromat-63X/1.4 M27 oil lens. 15-30µm z-stack images with a 0.6µm step-size were taken at 63x magnification. 60-120µm z-stack images with a 4µm step-size and single XY-plane snapshots were taken at 20x magnification. ST structures were identified as apically located multinucleated structures. vCTs were identified as E-cadherin-positive or integrin-⍺-6-positive mononucleated cells. vCT cell characteristics were measured using 63x z-stack confocal microscopy images on placental samples stained with anti-E-cadherin, phalloidin, and Hoechst. vCT cell length, vCT volume, vCT shape factor, relative nuclear positioning within vCT, and vCT apical volume was quantified using Volocity Imaging software (Quorum Technologies Inc., version 7.0.0). Relative positioning of the nucleus was calculated along the apical-basal axis by measuring the length between the center of the nucleus to the basement membrane and then dividing it by the total length of the cell (equation 1).

ST coverage and fusion capacity was measured using 20x single-plane confocal microscopy images on placental samples stained with anti-E-cadherin, phalloidin, and Hoechst. ST coverage was quantified using Volocity Imaging software by measuring the ratio of the cross-sectional area of the ST compared to the vCT (ST:vCT) (equation 2). Time dependent relative change in ST coverage was calculated as the relative ratio of ST:vCT coverage at 72hrs compared to 24hrs (equation 3). Relative fusion in ezrin-inhibitor-treated explants was normalized to vehicle treated explants.

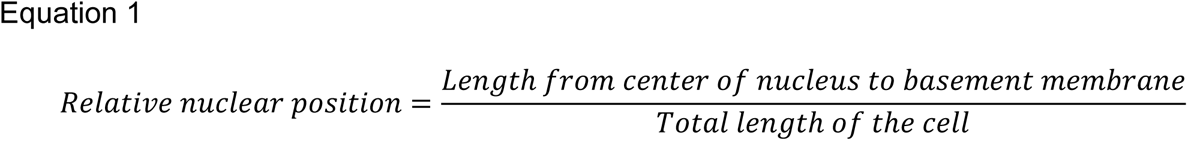

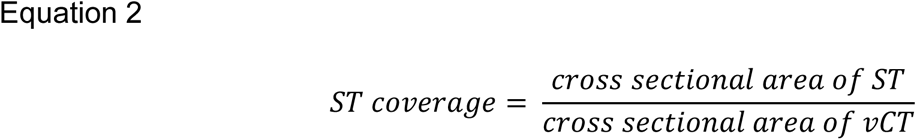

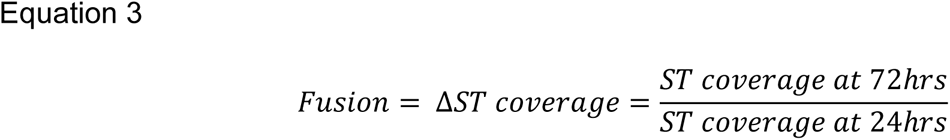

### Scanning electron microscopy (SEM)

At 24 hrs post-trypsinization, explants were collected, washed in PBS, and fixed in EM fixative (2.5% glutaraldehyde, 2% PFA in PBS). Tissue was then washed with PBS and treated with 1% osmium tetroxide in PBS for one hour. After washing, samples were dehydrated by incubation in a graded ethanol series and then in a graded series of increasing hexamethyldisilazane (HMDS) and decreasing ethanol concentrations. Samples were incubated in 100% HMDS and then airdried overnight. Tissue was then mounted onto standard aluminium stubs and sputter coated with Au/Pd using a Hummer 6.2 Sputter Coater (Anatech). Images were acquired using a Zeiss EVO 10 scanning electron microscope operating at 15kV and SmartSEM software (Zeiss, version 6.06).

### Transmission electron microscopy (TEM)

PBS washed and trimmed placental tissue was fixed in EM fixative (2.5% glutaraldehyde, 2% PFA in PBS) upon collection. Tissue was then washed with PBS and treated with 1% osmium tetroxide in PBS for one hour. After washing, tissue was then dehydrated by incubation in a graded ethanol series followed by incubation in 1:1 ethanol:Spurr resin (Electron Microscopy Sciences; Hatfield, PA, USA) overnight. The following day the tissue was embedded in 100% resin and cured overnight at 70°C. Using a Reichert-Jung Ultracut_E Ultramicrotome, the resin blocks were cut into 70-90nm sections. Sections were stained with uranyl acetate and lead citrate on 75 mesh grids. Images were acquired using a Philips/FEI (Morgagni) transmission electron microscope operating at 80kV with Gatan camera and Digital Micrograph software (version, 1.81.78).

### Reverse-transcriptase polymerase chain reaction (RT-PCR)

After 48 hrs of incubation (+/- 50µM ezrin inhibitor), explant cultured tissue was collected, rinsed in PBS, and then homogenized in TRIzol (Thermo Fisher Scientific) using a tissue lyser and RNA was extracted using TRIzol-chloroform extraction. RNA was isolated and purified using a PureLink RNA Mini Kit (Ambion; Carlsbad, CA, USA ; cat:12183025). Conversion into cDNA was performed using reverse transcription (iScript cDNA Synthesis Kit; Bio-Rad Laboratories, Mississauga, ON, Canada; cat:1708890) with 250ng RNA. Quantitative RT-PCR was performed using 3uL diluted cDNA (1:5) per reaction and Applied Biosystems SYBR Green PCR Master Mix (Thermo Fisher Scientific; cat:4309155) on a QuantStudio 3 Real-Time PCR System (Thermo Fisher Scientific). Primer sequences are indicated in Appendix Table 2. Relative change in mRNA expression was calculated using the 2-^ΔΔCT method to paired samples using the mean CT from *CYC1* and *TOP1* as housekeeping genes (Kaitu’u-Lino et al., 2014; Livak and Schmittgen, 2001; Shaha et al., 2022).

### Statistical analyses

Statistical analyses were conducted using GraphPad Prism (version 9.3.1) with a p<0.05 threshold for significance. Exact statistical analyses are within the figure legends. Statistical outliers were determined using a ROUT outlier analyses in GraphPad Prism. All graphs and representative images are representative of three to six biological replicates (placental tissue from different patients) and at least three technical replicates.

## Acknowledgements

We would like to thank all the tissue donors and the staff of Women’s Health Options Clinic without whom this work would not have been possible. We would also like to thank Dr. Kacie Norton and Arlene Oatway of the University of Alberta Biological Sciences Imaging Facility for their expert technical advice for the electron microscopy sample preparation and imaging.

## Data Availability

All relevant data can be found within the article and supplementary information.

## Diversity and Inclusion Statement

The authors W.D (she/her), S.Z.S (she/her), I.D (he/him), and K.P (she/her) have lived experience as racial and ethnic minorities in Canada intersecting with our respective diverse gender identities and sexual orientations. Ensuring equity, diversity, and inclusion in the field of developmental biology is more than moral obligation but also necessary to enrich existing and future knowledge and to promote excellence in research and healthcare. As a lab we are committed to integrating these diverse perspectives into our current understanding and providing equitable opportunities.

